# Genome-resolved viral ecology in a marine oxygen minimum zone (OMZ)

**DOI:** 10.1101/2020.07.22.215772

**Authors:** Dean Vik, Maria Consuelo Gazitúa, Christine L. Sun, Montserrat Aldunate, Margaret R. Mulholland, Osvaldo Ulloa, Matthew B. Sullivan

## Abstract

Oxygen minimum zones (OMZs) are critical to marine nitrogen cycling and global climate change. While OMZ microbial communities are relatively well-studied, little is known about their viruses. Here we assess the viral community ecology of 22 deeply sequenced viral metagenomes along a gradient of surface oxygenated to anoxic waters (< 0.02 μmol/L O_2_) in the Eastern Tropical South Pacific (ETSP) OMZ. We identified 46,127 viral populations (>5 kb), which augments the known viruses at this site by 10-fold. ETSP viral communities clustered into 6 groups that correspond to oceanographic features, with 3 clusters representing samples from suboxic to anoxic waters. Oxygen concentration was the predominant environmental feature driving viral community structure. Alpha and beta diversity of viral communities in the anoxic zone were lower than in surface waters, which parallels the low microbial diversity seen in other studies. Viruses were largely endemic as few (6% of viruses from this study) were found in at least another marine metagenome, and of those, most (77%) were restricted to other OMZs. Together these findings provide an ecological baseline for viral community structure, drivers and population variability in OMZs that will help future studies assess the role of viruses in these climate-critical environments.

**Originality-Significance Statement:** Marine oxygen minimum zones (OMZs) are unique and important ocean ecosystems where microbes drive climate-altering nutrient transformations. This study provides a baseline, deeply sequenced viral metagenomic dataset and reference viral genomes to assess ecological change and drivers across the oxygenated surface to de-oxygenated deep waters of the Eastern Tropical South Pacific (ETSP) OMZ. Community ecological assessment of the ETSP viromes reveals a relatively low diversity viral community with a high degree of endemic populations in the OMZ waters.

## Introduction

Oxygen deficient regions of the ocean play a vital role in regulating the ocean nitrogen budget and greenhouse gas emission (Wright *et al*., 2012). These low oxygen regions termed Oxygen Minimum Zones (OMZ) result from a combination of thermal stratification of the water column, low circulation, temperature- and wind-driven upwelling currents, and heterotrophic consumption of surface water primary production in deep water (Wyrtki, 1965; Kessler, 2006; Karstensen *et al*., 2008; Paulmier & Ruiz-Pino, 2009; Czeschel *et al*., 2011). Oxygen concentrations in OMZs can reach below the detection limit of a few nanomoles per liter in certain regions such as the Eastern Tropical South Pacific (ETSP) (Revsbech *et al*., 2009; Thamdrup *et al*., 2012; Ulloa *et al*., 2012), creating anoxic marine zones (AMZs). Problematically, OMZs have expanded over the last 50 years as a result of increased ocean stratification from rising surface water temperatures (Stramma *et al*., 2008, Wright *et al*., 2012, Ulloa *et al*., 2012). As OMZs expand, the metabolisms found therein – anaerobic ammonium oxidation (anammox) and denitrification – result in large-scale nitrogen loss through the increased production of N_2_ and the potent greenhouse gas N_2_O (Lam and Kuypers, 2011). Thus, the factors moderating microbe-mediated nutrient cycling in OMZs require rigorous examination to better understand the impact of OMZs on climatic trends.

The biological factors responsible for the development and maintenance of OMZs are almost exclusively a result of microbial activity (Zakem *et al*., 2019). As oxygen is removed from the environment by heterotrophs, other electron acceptors (such as nitrate and sulfate) that have progressively lower electron affinities and energy potentials are used. This results in a gradient of electron acceptors and redox chemistry across the depth gradient, referred to as the redoxycline. Due to the reduction in free energy, the biogeochemistry of OMZs is almost entirely dictated by the microbial metabolisms stratified along the redoxycline, rather than macrofaunal respiration (Hawley *et al*., 2014). This reduction in energy may be expected to be associated with a reduction in community size and diversity possibly due to a depletion in niche space (Rabosky, 2009; Beman & Carolan, 2013). However, major trends in microbial diversity across OMZs remain unclear, as diversity has been shown to either increase or decrease with the reduction of oxygen concentration (Stevens & Ulloa, 2008; Bryant *et al*., 2012). Nevertheless, it is plausible that diversity trends follow patterns in productivity, *i*.*e*., higher relative diversity in the surface chlorophyll maximum (SCM) and the deep chlorophyll maximum (DCM) (Chase & Leibold, 2002; Walsh *et al*., 2016), or the availability of niche space, which appears to peak at the interface between oxygenated and anoxic water, a transition environment where a wide array of metabolisms may persist (Bertagnolli & Stewart, 2018).

Importantly, viruses have also been shown to play major roles in both bottom-up and top-down mechanisms controlling microbial communities. In the surface oceans, viruses mediate microbial population dynamics and metabolism (Suttle, 2007; Hurwitz *et al*., 2013) through viral lysis, which in addition to protist grazing, results in microbial mortality rates that are proportional to growth rates at ∼1-2 day^-1^ (Ducklow & Hill, 1985; Cole *et al*., 1988; Suttle, 1994; Fuhrman & Noble, 1995). The result of this lysis is the redirection of fixed carbon away from macrofaunal production and into both microbial respiration and carbon export (Azam *et al*., 1983; Fuhrman, 1992; Guidi *et al*., 2016). In addition, viruses have been shown to encode host-derived auxiliary metabolic genes (AMGs), with a few notable examples of viruses likely associated with sulfur and nitrogen cycling (Roux *et al*., 2016; Ahlgren *et al*., 2019). Though data to date suggest that OMZ viruses are likely important, a foundational ecological perspective on viruses in these habitats is lacking.

To date, only two genomic studies have explored community dynamics of viruses in OMZ systems (Cassman *et al*., 2012; Roux *et al*., 2014). Both studies were relatively small in terms of samples collected, sequencing available, and viruses recovered, but both led to significant advances in our understanding of OMZ viruses. Specifically, Roux *et al*. (2014) focused on viruses that were recovered from 127 single cell amplified genomes from uncultivated SUP05 bacteria in a model OMZ ecosystem while Cassman *et al*. (2012) examined the diversity and size of viral communities in the ETSP OMZ region through metagenomics. Fortunately, sequencing costs and analytics has improved considerably since these initial studies, which warrants re-investigation of these viral communities. Here, we deeply sequence viral metagenomes from 22 seawater samples to provide 420-fold more sequencing data from these environments to establish genome-resolved datasets for exploring viral community and population ecology along oxygenated to anoxic marine waters of the ETSP OMZ.

## Results and Discussion

Samples were collected from six stations spanning a transect from coastal to pelagic waters in the ETSP OMZ region off the coast of Lima, Peru (**Figure 1A**, see Experimental Procedures). The depths at which samples were collected were determined by distinct oceanographic features (**Figure 1B, Table S1, Figure S1**). At most stations a sample was collected from the surface chlorophyll maximum, the upper oxycline, the upper OMZ (with or without a secondary deep chlorophyll maximum), and the core of the OMZ (**Figure 1**). Viral concentrates produced for each sample were sequenced with Illumina HiSeq 2000 to produce an average of 7.2 Gb bases per sample (**Table S2**). To provide context with other large studies, the Global Ocean Virome dataset had an average of 8.9 and 27.2 Gb per sample, on their first and second versions, respectively (Roux *et al*., 2016; Gregory *et al*., 2019). Reads were assembled into scaffolds, from which viruses were identified and then clustered into viral populations (see Experimental Procedures) that represent approximately species level viral taxonomy (Brum *et al*., 2015; Gregory *et al*., 2016; Gregory *et al*., 2019). In total, we recovered 46,127 viral populations of at least 5 kb in length and used these for all analyses in this study.

**Figure 1.**
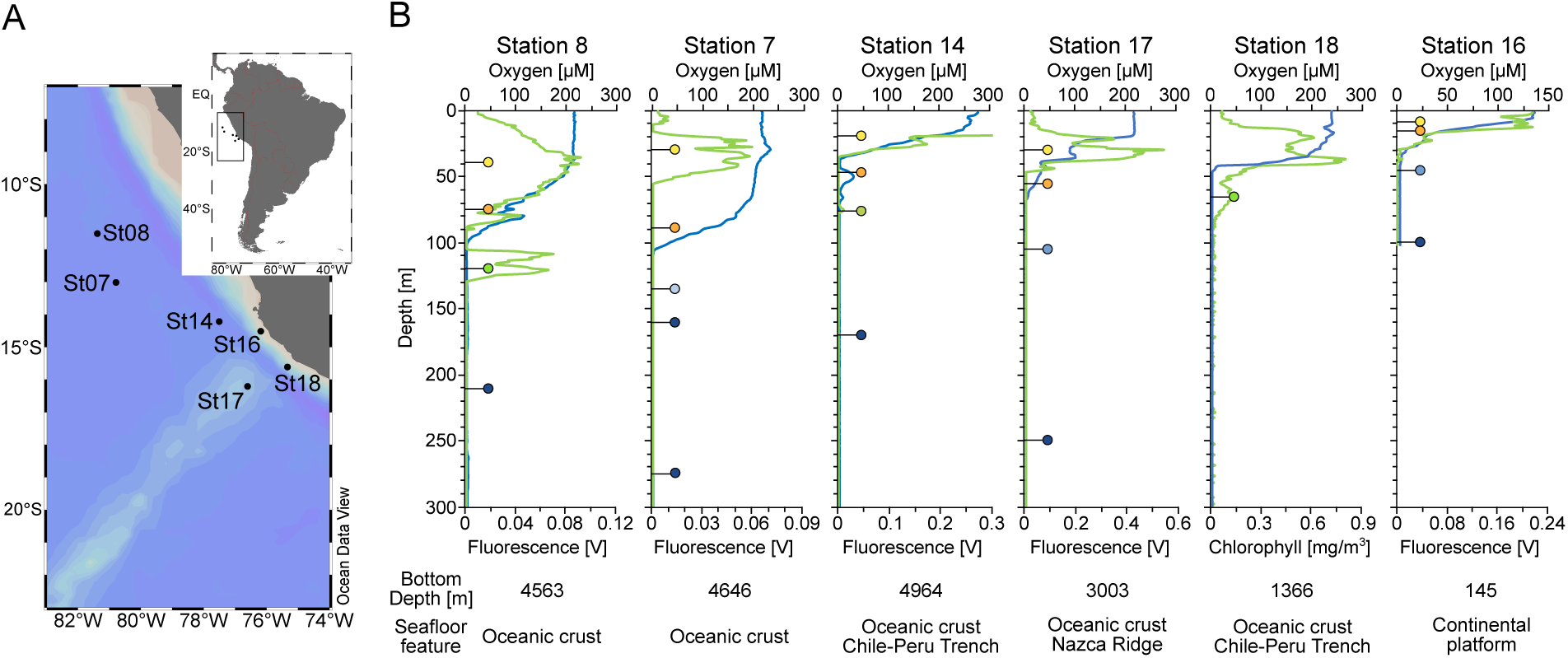
Map of the study area and vertical characterization of the sampling stations. **A**. Location of stations 7, 8, 14, 16, 17, and 18, off the coast of Peru in the ETSP OMZ. The map was created with Ocean Data View (http://odv.awi.de). **B**. Oxygen (solid blue line) and fluorescence/chlorophyll (solid green line) depth profiles from each station. Samples were collected at depths indicated by lines with the colored circles to depict the sample type: surface chlorophyll maximum in yellow, oxycline in orange, upper OMZ with deep chlorophyll maximum (DCM) in green and without DCM in light blue, and OMZ core in dark blue.

The relative abundances of the viral populations in each sample were calculated and normalized across all samples (see Experimental Procedures) and used as input for biogeographical and diversity analyses (**Table S3**). We hypothesized that viruses would cluster into distinct communities from the anoxic and oxic waters due to the environmental differences of these marine habitats (Bertagnolli & Stewart, 2018). This would be similar to previous studies that have shown OMZs have unique microbial communities, compared to more oxygenated surface waters (Madrid *et al*., 2001; Fuchs *et al*., 2005; Stevens & Ulloa, 2008; Wright *et al*., 2012). Overall, 6 main clusters were detected via hierarchical clustering (**Figure 2**), which is relatively consistent with those predicted from other statistics including a gap statistic (**Figure S2**) and an affinity propagation analysis (**Figure S3**). As expected, the anoxic waters of the OMZ were significantly distinct from the rest of the samples (**Figure 2**). Within the OMZ samples, there were 3 sub-clusters: samples from coastal OMZs (OMC_C cluster), samples from the upper OMZ with a DCM (uOMZd cluster), and samples from the remaining upper and core OMZ from pelagic waters (OMZ_P cluster) (see Experimental Procedures for cluster name information). Five of the samples from OMZ_P cluster also have relatively high concentrations of nitrite (**Table S1** and **Figure S1F**), and so represent an anoxic marine zone (AMZ) (Thamdrup *et al*., 2012; Ulloa *et al*., 2012). However, for the rest of the OMZ_P cluster samples, nitrite measurements are missing (see Experimental Procedures). Other clustering differences showed that pelagic surface waters differed from the oxycline and coastal surface waters. The main clusters support the conclusion that viral communities that exist in OMZs are relatively distinct from those communities found in the ocean surface.

**Figure 2.**
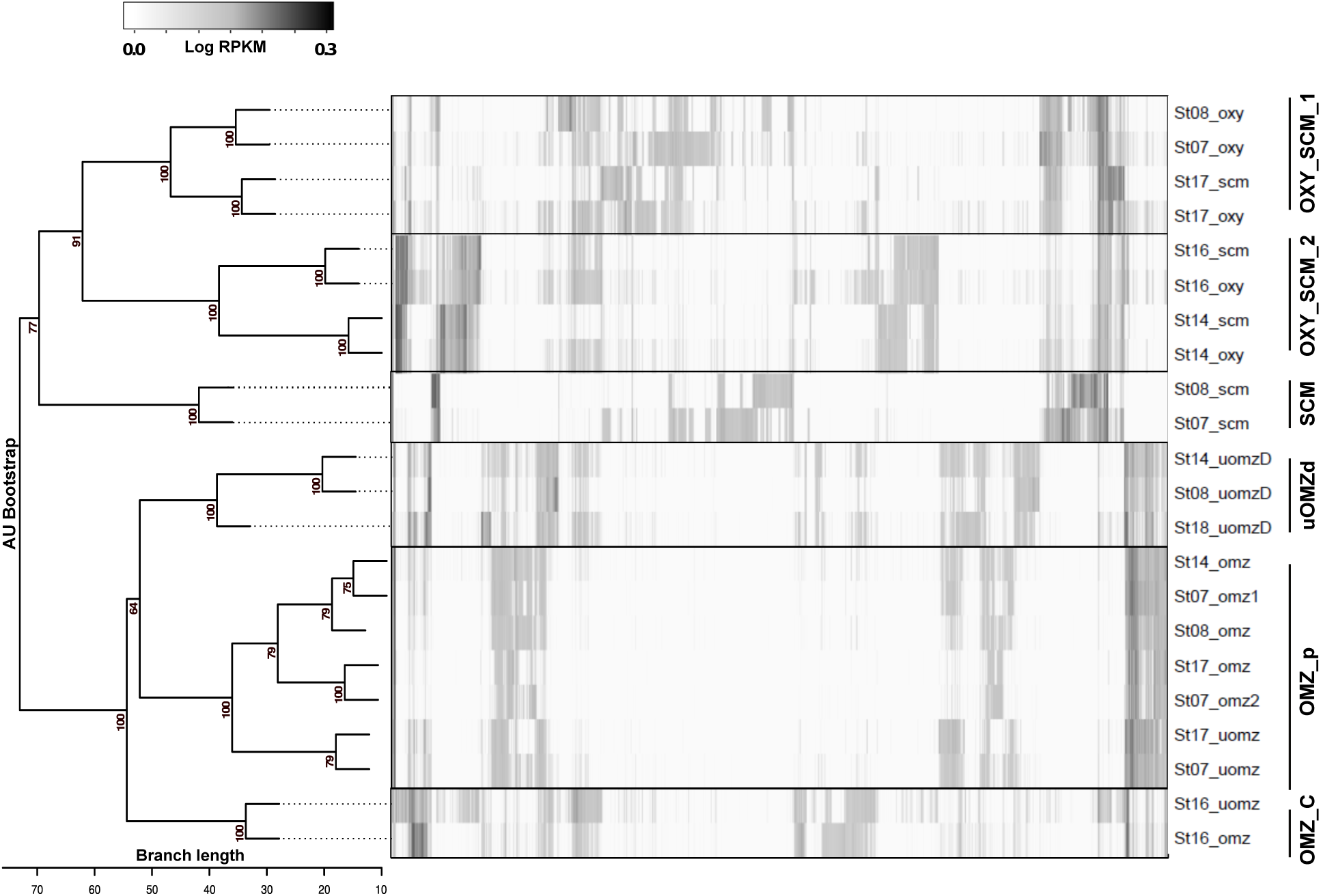
Hierarchical clustering of samples based on normalized relative abundances of viral populations revealed 6 clusters. Each row represents a different sample, labelled by station and sample type (scm = surface chlorophyll maximum, oxy = oxycline, uomzD = upper OMZ with deep chlorophyll maximum, uomz = upper OMZ without DCM, and omz = omz core). Each column represents a different viral population (≥ 5kb), where the normalized relative abundance values (log_10_ transformed) are shown in grayscale. RPKM means Reads Per Kilobase, per Million mapped reads. “Approximately unbiased” bootstrapping values are represented as a proportion of 100 permutations.

We next evaluated our viral communities with various diversity measures, including evenness, alpha diversity, and beta diversity (**Figure 3A, 3B, 3C**, respectively). The diversity metrics used were specifically selected to minimize the impact of varying sequencing depth on diversity estimations. In terms of the entire system, species accumulation analysis revealed only a 2% increase in species recovery in the last of the 22 randomly permuted sub-samples, indicating sequencing approached saturation (**Figure S4)**. Statistically robust species accumulation analysis was not possible at the individual community scale due to a lack of samples within a given habitat. Evenness did not significantly differ between samples from oxygenated regions and the OMZ (**Figure 3A**, Kruskal-Wallis p-value = 0.742). The evenness was nearly 1 in all cases (range 0.965-0.978, mean 0.974), which indicates that no community or sample cluster had a high relative proportion of dominant viral populations.

**Figure 3.**
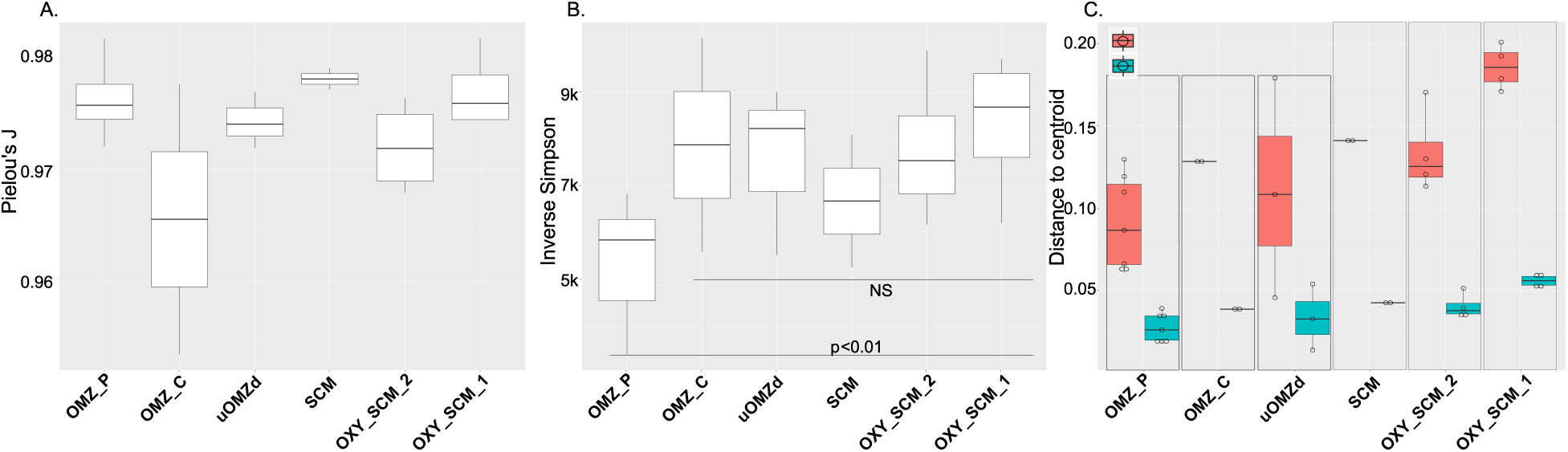
Diversity measures for the 6 main viral community clusters. **A**. Evenness across the 6 clusters. **B**. Alpha Diversity. **C**. Beta Diversity, where red corresponds to the beta diversity differences resulting from abundance while blue represents the beta diversity differences related to composition. The top and bottom of each box show the standard deviation while the line inside the box shows the mean. Points within each box represent the number of samples per community.

Alpha diversity, a measure of diversity within a community, was calculated (**Figure 3B**) using the inverse Simpson’s concentration (see Experimental Procedures), which facilitates a relatively unbiased comparison of alpha diversity across communities despite uneven sequencing depth (Haegeman *et al*., 2013). Alpha diversity did not differ between communities (Kruskal-Wallis p=NS) except for the OMZ_P cluster, which exhibited a significantly lower alpha diversity (Kruskal-Wallis p = 0.01). Beta diversity, a measure of the amount of the total diversity accounted for by a given community, was estimated (**Figure 3C**) via multivariate dispersion, which leverages distance-based ordination techniques to derive the average distance of all samples in a community from the community centroid (Anderson *et al*., 2006). Beta diversity was significantly lower in the OMZ regions than in the surface waters regarding both population composition (modified Gower’s distance log_10_ p = 0.005) and abundance (modified Gower’s distance log_2_, p = 0.006) (Anderson *et al*., 2006). The lower alpha and beta diversities for OMZ samples, consistent with a previous study (Cassman *et al*., 2012), indicate that niche space is reduced as energetics of the system decreases along the redoxycline. A similar trend has also been suggested for microbial community distributions in previous studies (Bryant *et al*., 2012; Beman & Carolan, 2013; Bertagnolli & Stewart, 2018).

Because the hierarchical clustering and diversity measures indicated that OMZ viral communities were distinct and relatively low in diversity, we next sought to identify the environmental features driving these patterns. To this end, we created ordination plots for the samples, as well as for the environmental features data (**Figure 4A, 4B**, *respectively*; stress plots **Figure S5**). Non-metric multidimensional scaling analysis (NMDS) with Bray-Curtis dissimilarity revealed that the 6 viral clusters (**Figure 2**) were distinct (**Figure 4A**, stress plots **Figure S5**). These findings were also supported statistically by ANOSIM (community R stat 0.855, p = 0.001 after 999 permutations) and by MRPP (distances within groups 0.528, between groups 0.872, overall 0.754, chance corrected within group agreement 0.351, p = 0.001). Together, these findings suggest ecologically distinct viral communities exist in samples within our dataset.

**Figure 4.**
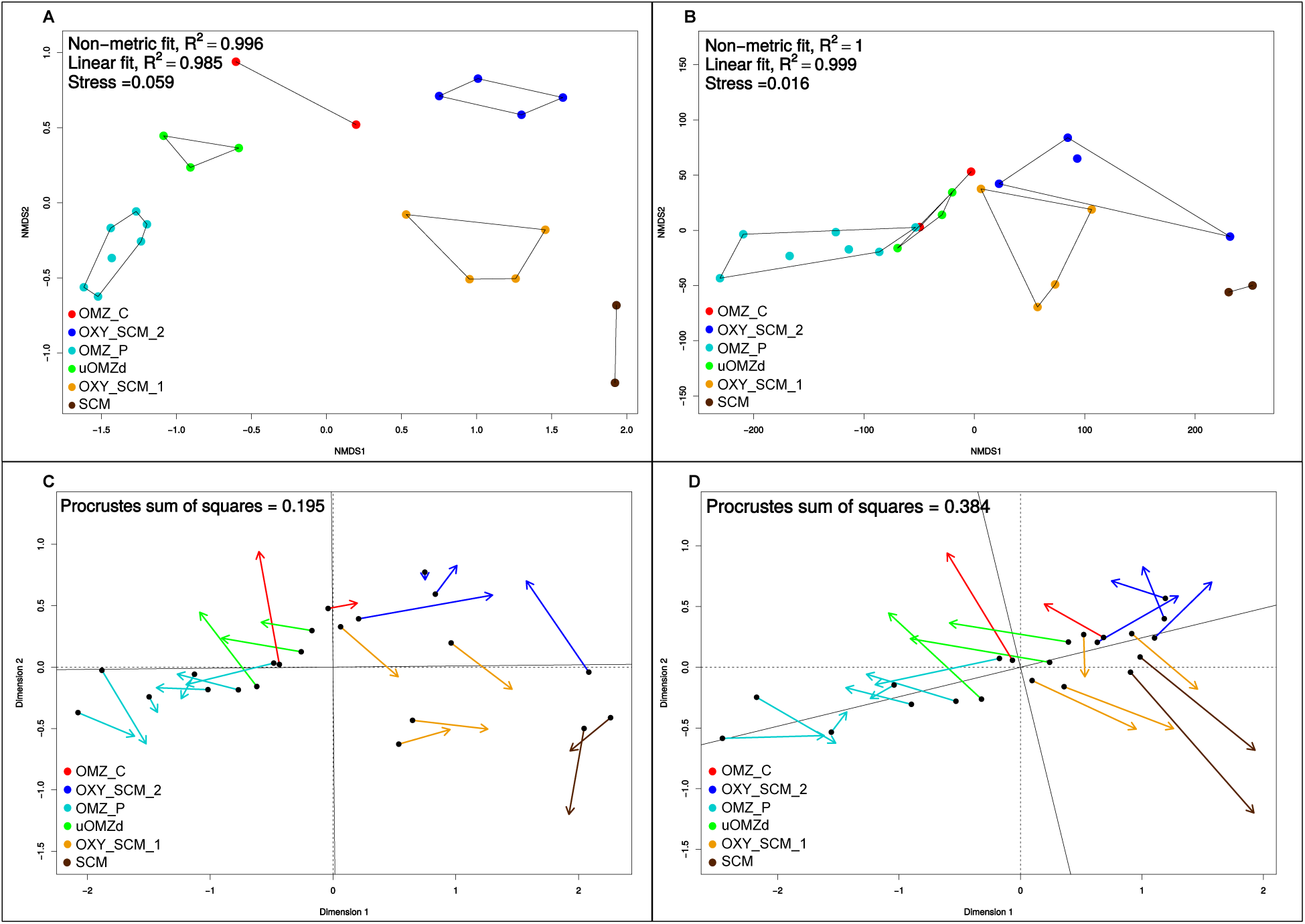
Environmental influence on the distribution of viral communities. **A**. Viral community ordination using NMDS and Bray-Curtis dissimilarities. Each of the 6 distinct communities are incorporated into the outlined clusters and color coded. **B**. Environmental feature ordination using NMDS and Euclidean distance. The colors are the same as in A. **C**. A comparison between the viral community ordination and environmental feature ordination where temperature has been removed from the latter dataset. Each arrow represents a sample’s spatial distance between ordinations, again with the same color coding. The statistical similarity between ordinations is represented as the Procrustes sum of squares where a lower value is more significant. D. The same comparison between the viral community ordination and environmental feature ordination, but with oxygen removed from the latter dataset.

In order to determine whether this separation between clusters could be explained by the measured environmental features, we compared ordinations based on viral populations (**Figure 4A**) and environmental features (**Figure 4B**). Similarities among the structures of these ordination plots and their underlying dissimilarity/distance matrices would indicate an environmental influence on the distribution of the viral communities. The structure of the viral community and environmental features ordination plots (Procrustes sum of squares 0.194, correlation 0.898, p = 0.001) (**Figure S6**) and trends in the dissimilarity/distance matrices (Mantels R = 0.675 p=0.001) were similar with the main structural difference being that the OMZ_P cluster was collapsed in the ordination created from the environmental features rather than separated into different OMZ sub groups. These results indicate that environmental features impact viral community distributions. While paired microbial community data were not available for comparison here, we posit, as done previously for surface ocean viral communities (Brum *et al*., 2013, Brum *et al*., 2015), that this reflects the environment associated biogeographical distribution of the resident microbial populations rather than a direct environmental impact on the viral populations.

Temperature and oxygen were the most descriptive gradients implicated in driving the biogeographical distributions of the viral communities (GAM and Pearson correlation; temperature p<0.0001, oxygen p<0.001). The co-variation of these two factors is expected in our system, as both decrease with depth, from surface to core OMZ waters (**Table S1, Figure S1A**, and **Figure S1B**). We addressed this co-variance by comparing the structure and agreement between NMDS ordinations of the environmental features and the viral community distributions again, but with the iterative removal of these parameters (removal of temperature in **Figure 4C**, removal of oxygen in **Figure 4D**) to determine which of these features was most important in retaining the similarity between these ordination analysis. The removal of temperature had an almost negligible impact on the relationship between the environmental features and the community distributions (Procrustes sum of squares 0.195, correlation 0.897, p = 0.001) indicating a relatively lower overall impact on the viral community structure. However, removal of oxygen reduced the relationship considerably (Procrustes sum of squares 0.385, correlation 0.784, p = 0.001), suggesting that, not surprisingly, oxygen was the most important driver of viral community composition (particularly the distinction between the communities found in the surface oxygenated water and the OMZ). Again, presumably this is due to the effect of oxygen on microbial populations rather than oxygen directly impacting the viruses.

Previous studies have indicated that OMZs have unique microbial and viral communities compared to the rest of the ocean (Madrid *et al*., 2001; Fuchs *et al*., 2005; Steven and Ulloa, 2008; Cassman *et al*., 2012; Wright *et al*., 2012). In order to determine to what extent the ETSP viral communities overlap with other oceanic viral communities, we evaluated whether our ETSP OMZ virus populations were among the ∼488K viral populations available in the Global Ocean Virome version 2.0 dataset (Gregory *et al*., 2019), and if so, assessed their biogeography. Using MMseq2, we identified viral populations from our study that shared 95% identity (over 50% of the ETSP query protein coding sequence) with the GOV2.0 populations (see Experimental Procedures). In total 2,763 of our 46,127 ETSP viral populations were also observed in the GOV2.0 dataset (**Figure 5A**), with about half (1,466) from OMZ samples (**Table S4**). Among these shared ETSP OMZ viruses, most (77%) were only found in other OMZ samples (O_2_ concentration below 10 µmol/L) (Helly & Levin, 2004) (**Table S4**). This shows that virtually all of our ETSP OMZ viral populations are endemic to OMZs, which is consistent with prior work (Cassman *et al*., 2012) where viral genotypes were evaluated (rather than viral populations) and where the geographical context was drastically reduced (only 4,552 viral genotypes were available for comparison as opposed to the 46,127 assessed here).

**Figure 5.**
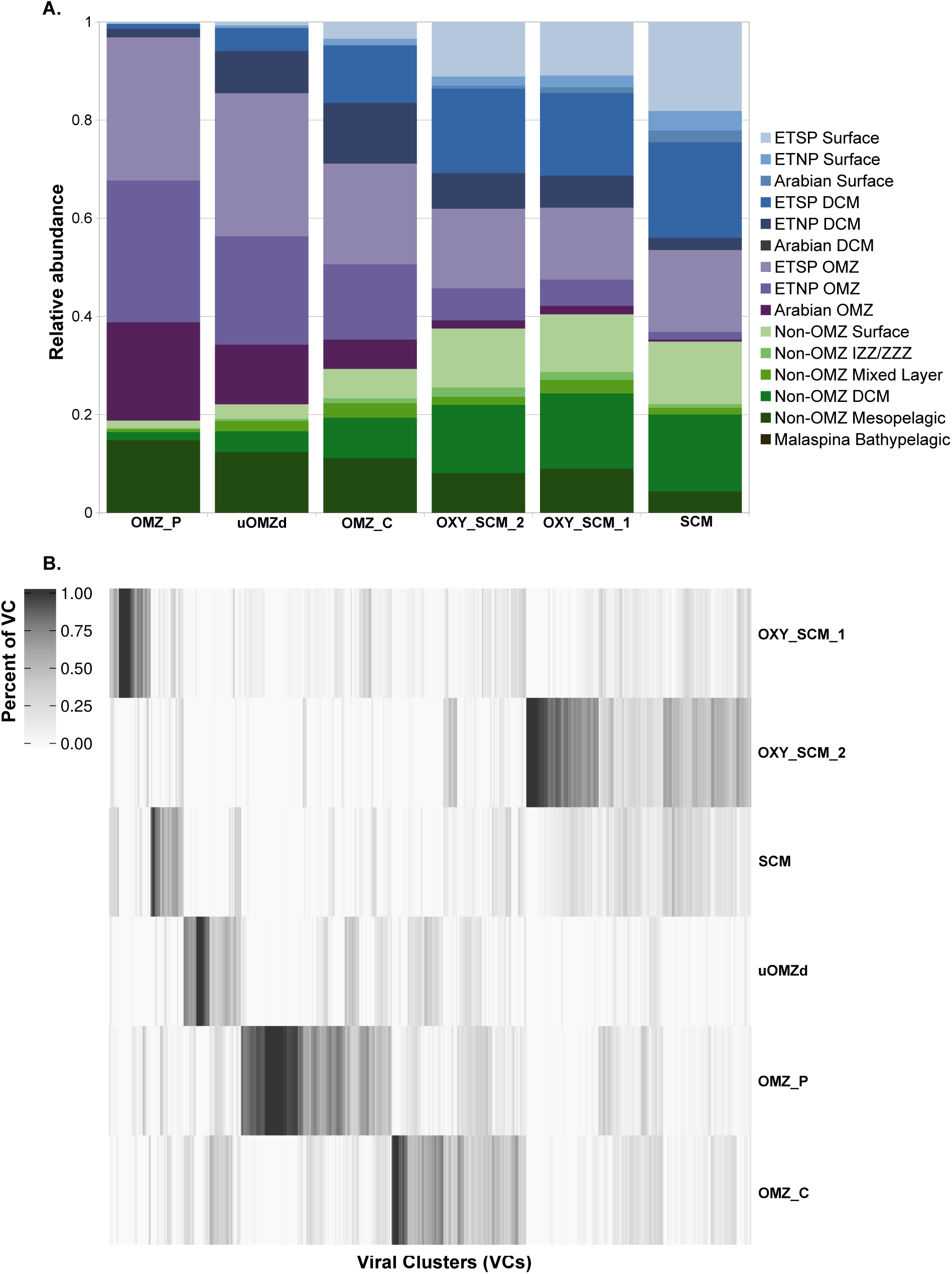
Sequence similarity to GOV2.0. **A**. ETSP sequence matches to GOV2.0 sequences, separated and color coded by GOV2.0 sample type. Each bar along the x-axis represents an ETSP community and each color along the y-axis represents the percent of relative abundance per GOV2.0 hit. Relative proportions within each community were calculated by summing relative abundances of each population within a GOV2.0 hit location, and then dividing that sum by the total sum of relative abundances with GOV2.0 hits (see Experimental Procedures). **B**. Distribution of viral clusters (viral genera) across each of the 6 communities. Each bar along the x-axis represents one viral cluster, colored by the percent of VC found in given community.

To further explore how the identified ETSP viral populations compared to known viruses in the RefSeq database, we used gene sharing networks where viral clusters (VCs) are approximately genus level taxonomy (Bolduc *et al*., 2017, Jang *et al*., 2019). With the sequences from viral RefSeq (v85) and the 10kb and larger viral populations in this study, these analyses clustered 10,632 viral populations into 1,465 VCs (**Figure 5B**), 4,020 viral populations into outliers (where populations were assigned to a VC but shared fewer similar proteins than the bulk of the cluster), and 482 viral populations into singletons (populations that did not cluster with any other sequences). Only 27 VCs included known reference viral sequences, which suggests that 98% (1438/1465) of the VCs derived from the ETSP OMZ dataset likely represent novel viral genera. If true, this is a 5-fold expansion of viral genus sequence space recovered from our analysis, as compared to RefSeq. Within the ETSP, 28% of the VCs identified in the OMZ sample were not present in any of the surface or oxycline samples further suggesting that the OMZ sample is distinct from the oxic habitats

Finally, we sought to use read mapping against our expanded dataset of ETSP viral populations to provide a very gross metric of population stability in these systems as assessed against the previous shallow viral metagenomic sequencing (Cassman *et al*. 2012). Less than 3% of the reads from *Cassman et al*. recruited to the ETSP viral populations, which corresponded to either as little as 1 or as much as 698 ETSP viral populations (using conservative vs permissive coverage cut-offs, see Experimental Procedures) being present in the prior dataset. This may represent high turnover in viral populations, but the inference does suffer from ascertainment bias due to the minimal sequencing available in the prior study.

## Conclusions

OMZs have been expanding over the last 50 years as a result of rapidly escalating anthropogenic carbon dioxide emissions increasing atmospheric temperatures, which in turn has increased ocean temperatures and stratification – features that select for OMZ formation and expansion (Schmidtko *et al*. 2017). With these ocean changes, and the fact that the oceans are a major carbon dioxide sink where microbes control that carbon’s fate, it becomes critical to understand how microorganisms will respond to and impact such changes (Cavicchioli *et al*., 2019). Viruses that infect these microbes also become important to understand. In this study, we present the largest survey of viruses from an OMZ – 46,127 unique viral populations across 6 stations at the ETSP OMZ (**Figure 1**). In ETSP, OMZ viral communities were distinct and relatively low in diversity compared to oxygenated, surface waters (**Figure 2**; **Figure 3)**, with oxygen as the most important driver of viral community composition (**Figure 4D**). These viruses are more similar to viruses from other OMZs and are novel (**Figure 5A, Figure 5B**). This is congruent with previous studies that have shown OMZs have unique and low diversity microbial communities, compared to the rest of the ocean (Madrid *et al*., 2001; Fuchs *et al*., 2005; Wright *et al*., 2012), which may result from the reduced redox potential of the prevalent electron acceptors in OMZs.

Though a large study, limitations are as follows. First, we cannot link the viruses to their microbial hosts because there is a lack of metagenomic samples from which we could construct metagenomically assembled genomes (MAGs), and such co-sampled MAGs improve virus-host linkages typically 5-fold or more (Emerson *et al*, 2018). Though AMGs are important in the surface oceans (reviewed in Rosenwasser *et al*., 2016, Hurwitz & U’Ren, 2016), they were not studied here as they are the focus of a parallel study from the same dataset that revealed viral genomes that contain AMGs associated with the denitrification, nitrification, and other nitrogen cycle processes, suggesting that these OMZ viruses influence the nitrogen cycle (Gazitua *et al, submitted*). Future work in OMZs should be enabled by our current findings and the vast sequence database of reference virus genomes that will empower a new generation of researchers to evaluate viral roles in modulating microbial population dynamics and biogeochemical cycling climate-critical OMZs as they expand due to climate change.

## Acknowledgements

We thank Sullivan Lab members and Heather Maughan for comments on the manuscript, and the crew of the *R/V Atlantis* for the sampling opportunity and support at sea. This work was funded in part by awards from NSF Biological Oceanography to MRM (#1356056), from the Agouron Institute to OU and MBS, a Gordon and Betty Moore Foundation Investigator Award (#3790) and NSF Biological Oceanography Awards (#0940390, #1536989) to MBS.

## Experimental Procedures

### Sample collection

On December 31, 2014 – January 22, 2015, six stations spanning a transect from coastal to pelagic waters in the ETSP OMZ region (off the coast of Peru) were sampled during the cruise AT-2626 aboard the *R/V Atlantis*. Volumes of 20 liters were collected using a pump profiling system (PPS), equipped with a Seabird SBE 25 Conductivity Temperature Depth (CTD), a WET Labs ECO-AFL/FL fluorometer, a Seabird SBE 43 dissolved oxygen sensor and a STOX sensor for nanomolar scale measurements of oxygen concentrations (detection limit of of 1-10 nmol L^−1^ O_2_). Oxygen detection limits using this sensor was about 0.02 μmol/L. High nitrite concentrations are found in waters with <50 nM of oxygen (Thamdrup *et al*., 2012). However, nitrite values in our sampling were only available for 3 samples (**Table S1**). In the cases where nitrite values were not obtained for a sample, nitrite values from adjacent depths (±10m) were used if available.

Concentrations of dissolved oxygen, nitrite, and other metadata can be found in **Table S1**. Sampling depths were selected according to variation in oxygen and chlorophyll concentrations, such as the surface chlorophyll maximum, the suboxic upper oxycline, the anoxic upper OMZ (with or without a deep chlorophyll maximum), and the core of the OMZ (**Figure 1, Table S1**). Samples for nitrite were filtered using a 0.2 µm cartridge filter. Filtrate was collected into sterile Falcon™tubes and stored upright at - 20°C until analysis. Nitrite concentrations were measured using an Astoria-Pacific autoanalyzer and standard colorimetric methods (Parsons *et al*., 1984), with a limit of detection (LOD) of 0.02 µM NO_2_ ^-^, (3σ, n = 7) (Selden *et al*., *submitted*).

Viral particles of the 22 samples were concentrated from the filtrate by iron chloride flocculation (John *et al*., 2011; Duhaime & Sullivan, 2012). Viral concentrates were then collected on a 1.0μm, 142mm, polycarbonate (PC) membrane (GE Water and Process Technologies, Trevose, PA, USA; Cat. #K10CP14220) and stored at 4 °C. The viral-iron precipitates were resuspended overnight in ascorbic-EDTA buffer (0.1 M EDTA, 0.2 M Mgcl, 0.2 M ascorbic acid, pH 6.0), rotating in the dark at 4°C. DNaseI at 100U ml^-1^ concentration was added to the final viral concentrate to remove any free DNA (Hurwitz *et al*., 2013). Viral DNA was then extracted using a Wizard DNA purification kit (Promega) with 1 ml resin to 0.5 ml sample. Samples yielding more than 1μg DNA (7 out of 22) were further purified using CsCl buoyant density gradients (Hurwitz *et al*., 2013). Viral contigs detected in the CsCl purified samples were retained only if they clustered into populations with viruses from the non-purified samples and became the representative contig of the population (the longest contig) (see *Assembly and processing*). Ecological analyses were then performed using the only 22 DNase-purified samples and representative contigs from all samples. DNA samples were submitted to JGI for library preparation and Nextera sequencing on an Illumina Hiseq 2000.

### Assembly and processing

Data processing and metagenomic analyses were performed using high-memory computer nodes from the Ohio State Supercomputer Center (Ohio State Supercomputer Center). Trimmomatic version 0.33 was used to remove Nextera adapters, to split reads into paired and unpaired groups, and to trim reads with low quality regions below a Phred score threshold of 15, using a sliding window of 4 bases (Bolger *et al*., 2014). Reads from each sample, with or without the CsCl purification step, were then assembled with Spades version 3.11.1, using the --meta option with paired end reads and the --sc and --careful options with unpaired reads, both with kmers of 21, 33, and 55 bases (Nurk *et al*., 2017). The resulting scaffolds were then clustered into population scale groups at 95% ANI over 80% of the shorter sequence using an in-house wrapper script for nucmer, run with default settings (Kurtz *et al*., 2004; Brum *et al*., 2015).

### Viral identification

Population contigs larger than 5 kb were processed with the viral identification tools VirSorter and Virfinder (Roux *et al*., 2015; Ren *et al*., 2017), and CAT (Cambuy *et al*., 2016), based on the steps described in Gregory *et al*. (2019). Populations with VirSorter categories 1 or 2, or with a VirFinder score ≥ 0.9 and a p-value < 0.05 were considered to be viral, as well those with VirSorter categories 3 to 6 and a VirFinder score >= 0.7 and a p-value < 0.05. Contigs with VirSorter categories 4 and 5 and a VirFinder score < 0.7 were manually curated to check if they were misannotated as prophages. If so, they were re-assigned to categories 1 or 2, respectively, and considered viral. Contigs with a VirFinder score between 0.7 and 0.9 (p-value < 0.05), without a VirSorter category, were run through CAT and those with < 60% of the genome classified as bacterial, archaeal, or eukaryotic (based on an average gene size of 1000) were considered viral. **Table S5** shows the VirSorter, VirFinder, and CAT assignments for each viral population.

### Viral relative abundances

In order to determine the relative abundance of each viral population larger than 5kb, the final viral populations were concatenated and then used as a database to recruit the quality trimmed reads using a custom wrapper script for bowtie2, which automatically determines groupings of paired and unpaired reads (Langmead & Salzberg, 2012). The resulting coverage files were then converted into a relative abundance table with the per population coverages using a custom wrapper script for BamM (https://github.com/ecogenomics/BamM). Coverages were calculated using the tpmean algorithm and adjusted coverages were calculated based on the coverage of each viral population per Gb of metagenome sequenced. Relative coverages were only reported if more than 75% of the population had at least 5x coverage, with at least 90% identity over 90% of the read. For reference, **Figure S7** shows the sequencing depth and number of reads that mapped to viruses for each sample.

In order to determine the fraction of the ETSP viral population that was also identified in the Cassman *et al*. (2012) study, the high quality, trimmed reads from Cassman *et al*. 2012 were downloaded from MGrast and recruited to the ETSP populations as described above for the abundance estimates. Due to a low number of ETSP viral populations being recovered with the stringent coverage threshold above, we eliminated the viral population coverage threshold to allow for more permissive read recruitment with the reads from Cassman *et al*. (2012).

### Cluster identification

Clusters were inferred using a combination of affinity propagation using the R function APCluster with options negDistMat(r=2) (Frey & Dueck, 2007; Bodenhofer *et al*., 2011) and a gap statistic using the R function clusGap with options kmeans, 10, and B = 100, (Tibshirani *et al*., 2001), resulting in an estimation of between 5 and 8 statistically supported groups. The relative abundances of the viral populations were then used in multiple permutations of a hierarchical clustering analysis with minkowski distances (p=2) to identify an approximation of the viral communities (Suzuki & Shimodaira, 2006). Viromes clustered with an approximately unbiased bootstrap value of 100% were considered viral communities. Viral population distributions among the viral communities were visualized with a heatmap plotted using the R package heatmap3. Bray Curtis distances were then plotted in a similar fashion to further validate the observed clustering patterns (Bray & Curtis, 1957).

The names of the 6 clusters in **Figure 2** were generated to be as descriptive as possible, using similar abbreviations from **Figure 1**. The clusters ‘OXY_SCM_1’ and ‘OXY_SCM_2’ denote clusters consisting of samples from oxycline and surface chlorophyll maximum. The ‘SCM’ cluster has samples from the surface chlorophyll maximum only. The ‘OMZ_C’ cluster has OMZ samples from the coastal St16, the ‘uOMZD’ cluster has samples from the upper OMZ with deep chlorophyll maximum, and the ‘OMZ_P’ has samples from the pelagic OMZ core.

### Comparison with environmental features

Distances within and between viral communities, as defined by the hierarchical clustering, were evaluated using nonmetric multidimensional scaling with Bray Curtis distances and 999 permutations or until convergence using the R package vegan and the function metaMDS (Field *et al*., 1982; Oksanen *et al*., 2018). The statistical significance of the viral community groups was then validated by comparing the within community and between community distances with MRPP and ANOSIM (Mielke *et al*., 1976; Clarke, 1993). Standardized Z-score and raw environmental feature measurements were then correlated with the viral community ordination using maximum linear and GAM non-linear algorithms using the R package envfit and odrisurf respectively (Clarke & Ainsworth, 1993). The environmental features that were used for these correlations are found in **Table S1**. Note that nitrite values were not used because they were only available from 3 samples directly (other samples had nitrite values taken from adjacent depths).

Known co-correlations between significant environmental features were then addressed by comparing distances between samples according to the relative abundances of the viral populations and the measured environmental features (Sunagawa *et al*., 2015). The standardized and raw environmental features were represented in ordination space using NMDS and Manhattan distances with 999 permutations until convergence (Field *et al*., 1982). Relationships between the standardized or raw environmental features and viral community ordinations were then evaluated, with and without the removal of a specific environmental feature of interest, using a Procrustes analysis and Mantel test (Mantel, 1967; Jackson, 1995). Analyses conducted with the standardized and raw environmental features were congruent, but more easily interpreted with the raw environmental features, and thus, results from the raw numbers are reported in the main text.

### Alpha diversity, beta diversity, and evenness

Diversity estimates were based on the relative abundance tables generated via read recruitment. Evenness, as a measure of the relative similarity among population abundances within a community, was calculated manually using equation H/ln(S) where H is the Shannon Wiener diversity index per community, calculated with the vegan diversity application in R using the option index = “shannon”, and S is the observed species richness of the community, calculated using the vegan application specnumber in R (Shannon, 1948; Pielou, 1966). Simpson concentration indices were calculated per viral community using the R package vegan and the application diversity with the option index = “invsimpson” (Simpson, 1949). Alpha diversities were represented as the inverse simpson concentration in order to facilitate the representation of statistically significant differences between communities (Jost, 2006) and to mitigate the uncertainties in diversity estimates due to variations in sampling effort (Haegeman *et al*., 2013).

We then compared beta-diversity among the 6 communities as a measure of the amount of the total diversity within a system accounted for by a given habitat, using a multivariate dispersion analysis. This approach facilitates attributing the observed diversity to only population composition or to population composition and abundance based on modified Gower distances (Anderson *et al*., 2006). Raw normalized relative abundance tables were first log transformed using the R application “decostand” and options method = “log” and logbase = 2 or 10. Distances were then calculated using the vegan application “vegdist” and the options method = “altGower” in R. The multivariate dispersion analysis was then performed on these distance matrices using the vegan application “betadisper”, “anova”, and “permutest” in R with defined groups from the hierarchical clustering analysis above.

### Endemism within ETSP

Viral populations were clustered into approximately genus level taxonomic groups using the network analytic vConTACT2 (Jang *et al*., 2019) in order to determine the relative proportion of each viral genus found within a community or shared across ETSP OMZ communities. Viral ORFs were first predicted and translated from the viral populations larger than 10 kb using Prodigal version 2.6.3 with the -p meta option (Hyatt *et al*., 2010). These predicted ORFs were then used to cluster the 10 kb populations amongst themselves and with viral Refseq version 85 using vConTACT2 with default parameters. Specific viral genera in each community were evident from the resulting network, so the relative abundance of each genus was determined by summing the relative abundances of the viral populations included in each genus. Genus abundance data were then tabulated and visualized using the R packages ggplots (Wickham, 2016).

Sequence comparisons were then used to determine the amount of ETSP viral populations larger than 5 kb that were identified in other regions of the ocean. MMseq2 using the easy-search command and with a 95% identity over 50% of the query protein coding sequence was used to compare the ETSP viruses with the 488k viral populations identified in the GOV2.0 database (Hauser *et al*., 2016; Gregory *et al*., 2019) (**Table S4**). The abundance and distribution of each GOV2.0 population identified was then evaluated to determine the habitats in which these populations were found. A stacked bar chart was then created to show the proportional abundance of each ETSP population with significant similarity to a population in GOV2.0, and the habitat wherein each GOV2.0 population was identified.

### Data availability

All high-quality reads and assembled contigs are available on iVirus (CyVerse, https://doi.org/10.25739/mmj5-kt58). Requests for further information should be directed to MBS at.

## Table and Figure Legends

**Table S1. Metadata for the 22 samples collected and sequenced**. Location and measurements of the selected environmental features sampled per site and depth. Samples are organized and labeled according to their respective cluster from the hierarchical clustering of the viral population abundances.

**Table S2. Sequencing information for each sample**. For each of the samples used in this study, this table lists the number of raw read, number of reads following quality control, the number of viral populations identified in each sample, and the number of reads that map to viral populations.

**Table S3. Relative abundances of the viral populations in each sample**. The abundances were normalized across samples via number of viral sequencing reads and by the length of each sequence.

**Table S4. Comparison of viruses from ETSP and GOV2**. Statistics for the ETSP viral populations with significant protein coding sequence similarity with GOV2.0, according to MMseq2 “easy-search”. Only sequences sharing 95% identity across 50% of the sequence are reported.

**Table S5. Categorization of viral populations**. For each of the 46,127 viral populations, this table contains the VirSorter, VirFinder, and CAT assignments.

**Figure S1. Vertical distribution of oxygen, temperature and salinity of the 6 sampled stations**. Oxygen, temperature and salinity depth profiles of the first 300 meters of each station. Colored circles indicate the depths where each of the 22 samples were collected: surface chlorophyll maximum in yellow (“scm”), oxycline in orange (“oxy”), upper OMZ with deep chlorophyll maximum (DCM) in green (“uomzD”) and without DCM in light blue (“uomz”), and OMZ core in dark blue (“omz”).

**Figure S2. Gap statistic for the number of significant sample clusters**. The cluster size, in number of samples, is where the observed within cluster distance is the smallest and yields the highest “gap” between expected within group distances. This was calculated using a null model, and observed within group distances. Here, clusters were derived by kmeans with 100 bootstraps and a maximize cluster size of 10.

**Figure S3. Affinity propagation analysis**. Clustering of samples according to negative squared Euclidean distances, using default input and exemplar preferences. The lighter yellow color corresponds with a higher similarity score, and each cluster is represented in the dendrograms with color coded bars.

**Figure S4. Species accumulation curve**. The number of species (viral populations) identified by 100 random sub-samplings of each of the pooled 22 samples across sampling depth and stations. Species richness estimations were computed using the jacknife2 estimator.

**Figure S5. NMDS stress plots of viral communities and environmental features**. A comparison of the fit of the distances displayed in the NMDS ordination plot with the true Bray Curtis dissimilarities between each station.

**Figure S6. Ordination plot of environmental features. Procrustes comparison of the environmental features and viral populations**. An alignment of the NMDS ordination plot created for the viral populations (Bray-Curtis dissimilarity) and environmental features (Euclidean distance). No environmental features were removed from this comparison. Procrustes sum of squares is 0.194.

**Figure S7. Sequencing depth**. Plot comparing the sequencing depth (gray bars), in terms of the number of post-quality control paired end reads, to the number of reads from that recruited to the identified viral populations, pooled from all samples (black bars).

