## Supplementary figures and images for "Genome-resolved viral ecology in a marine oxygen minimum zone (OMZ)"

### Figure S1

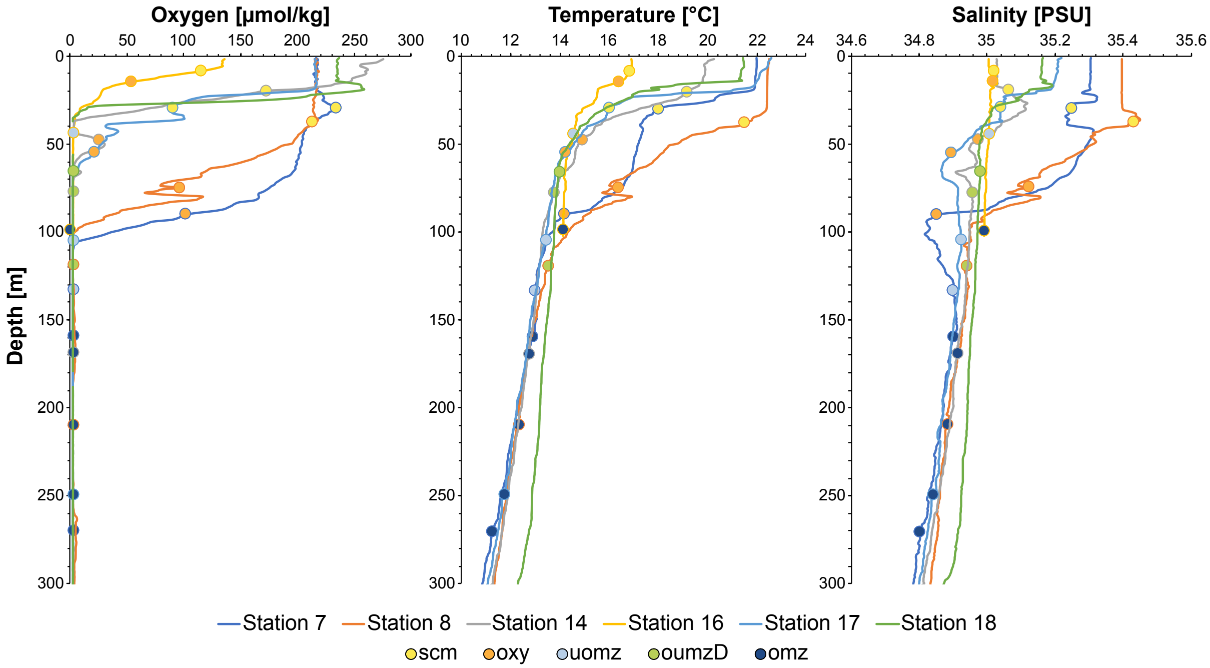

### Figure S2

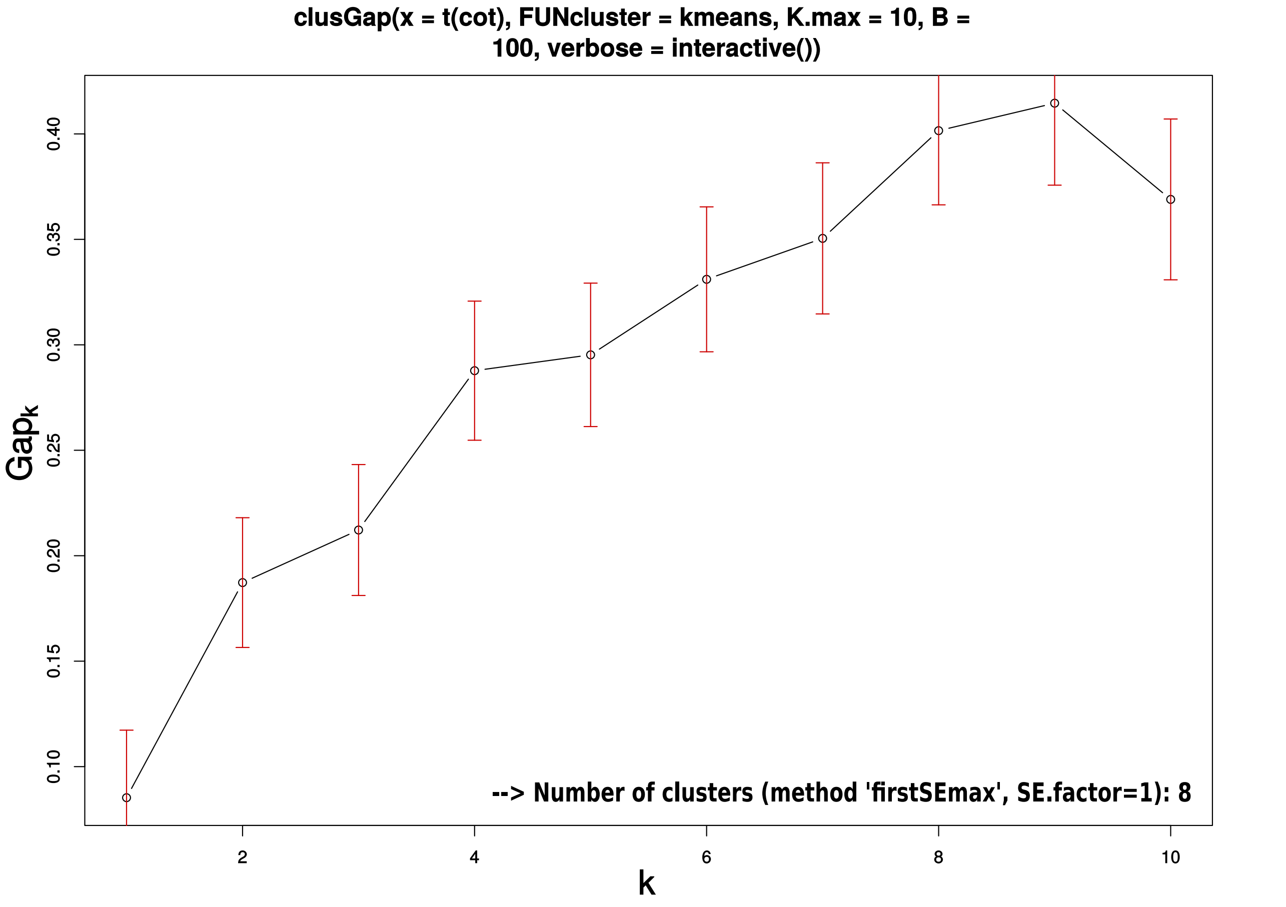

### Figure S4

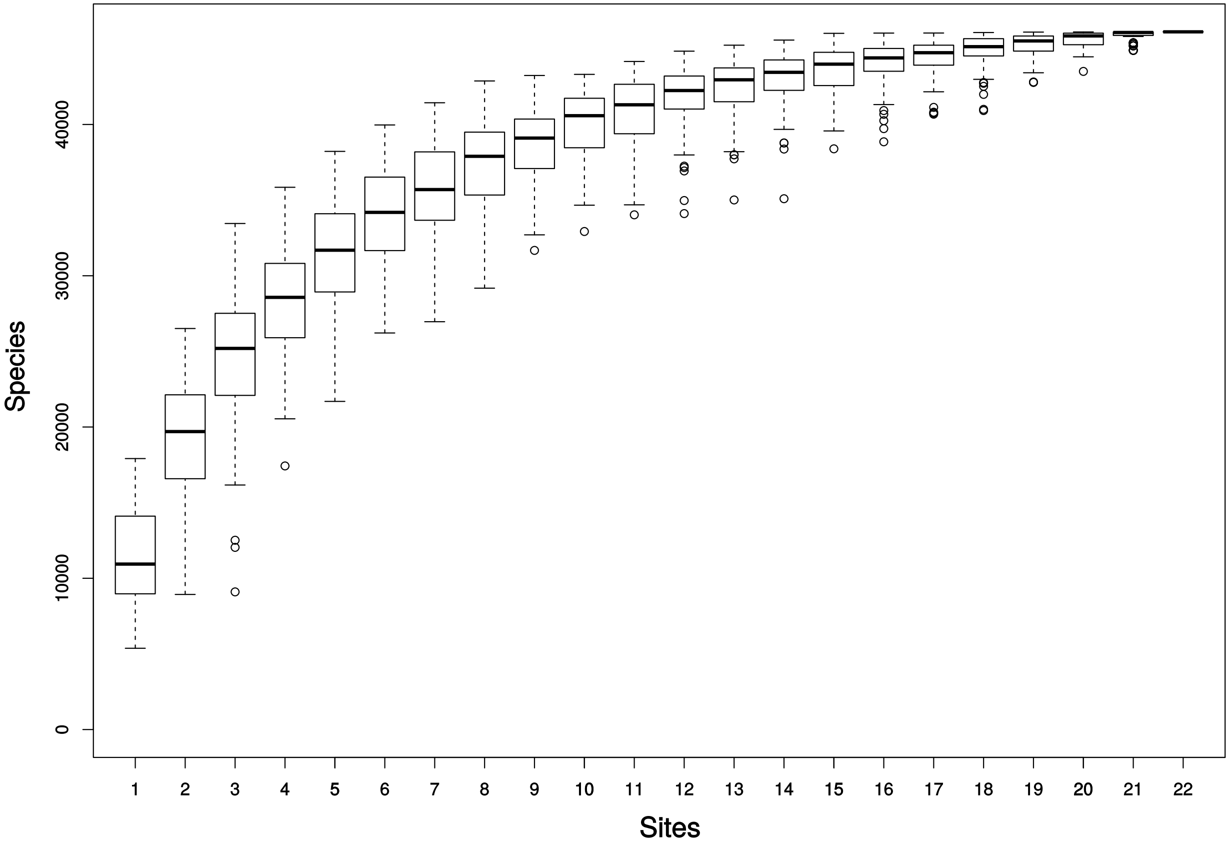

### Figure S5

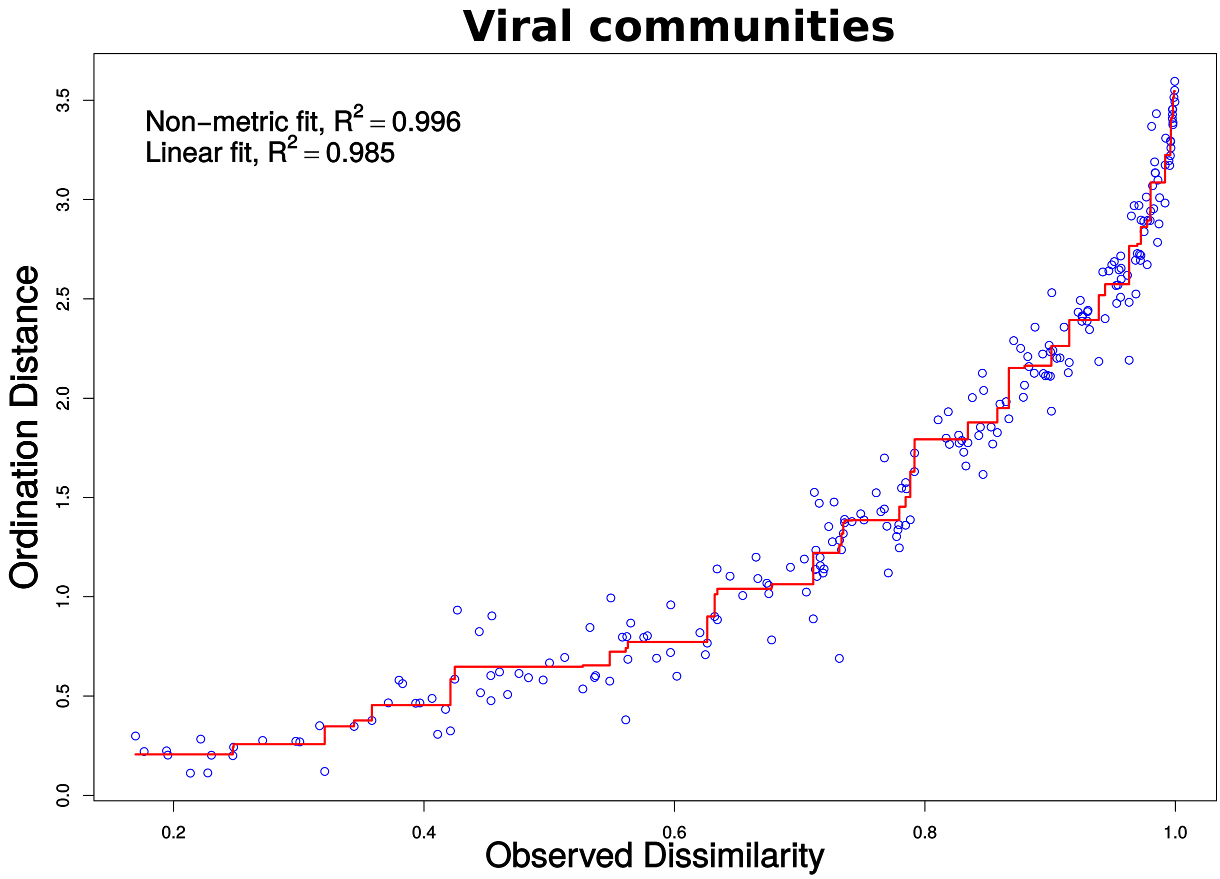

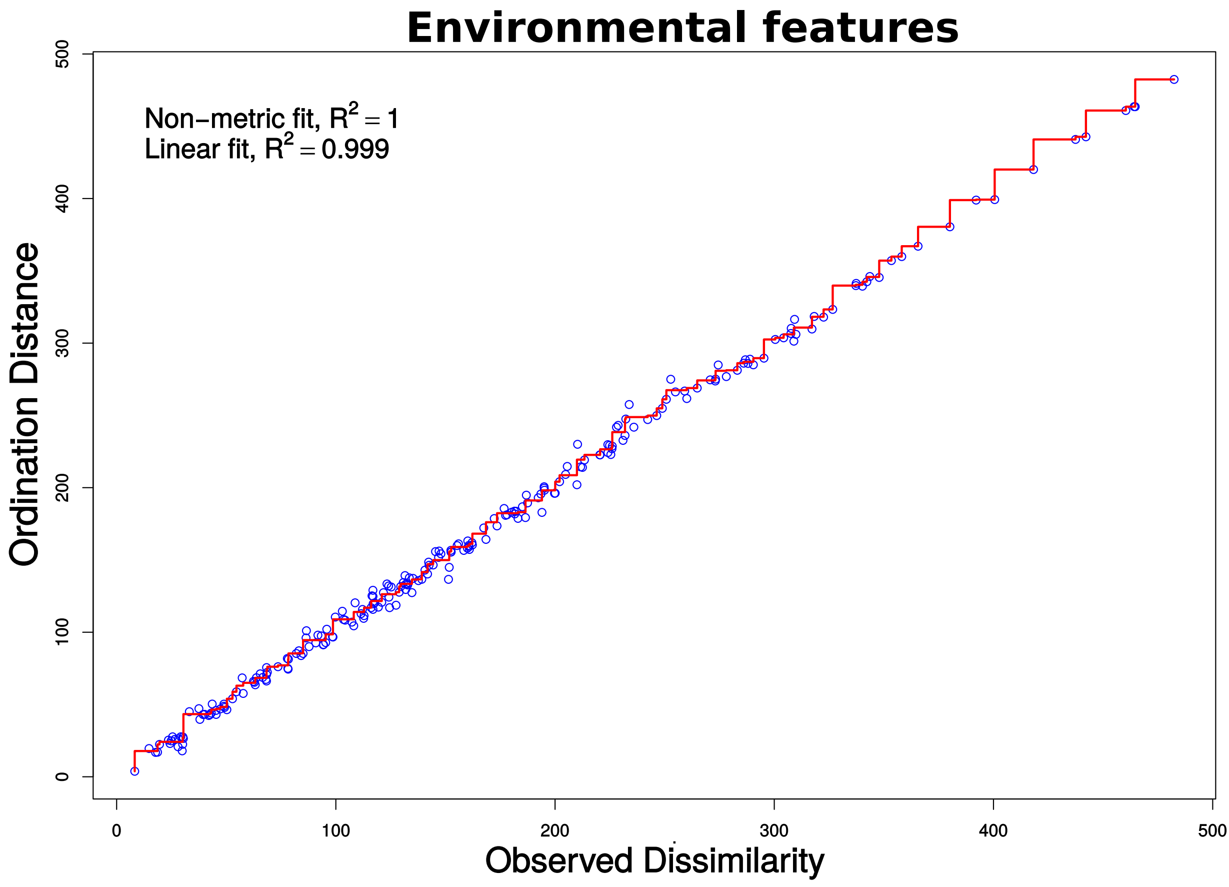

### Figures S6

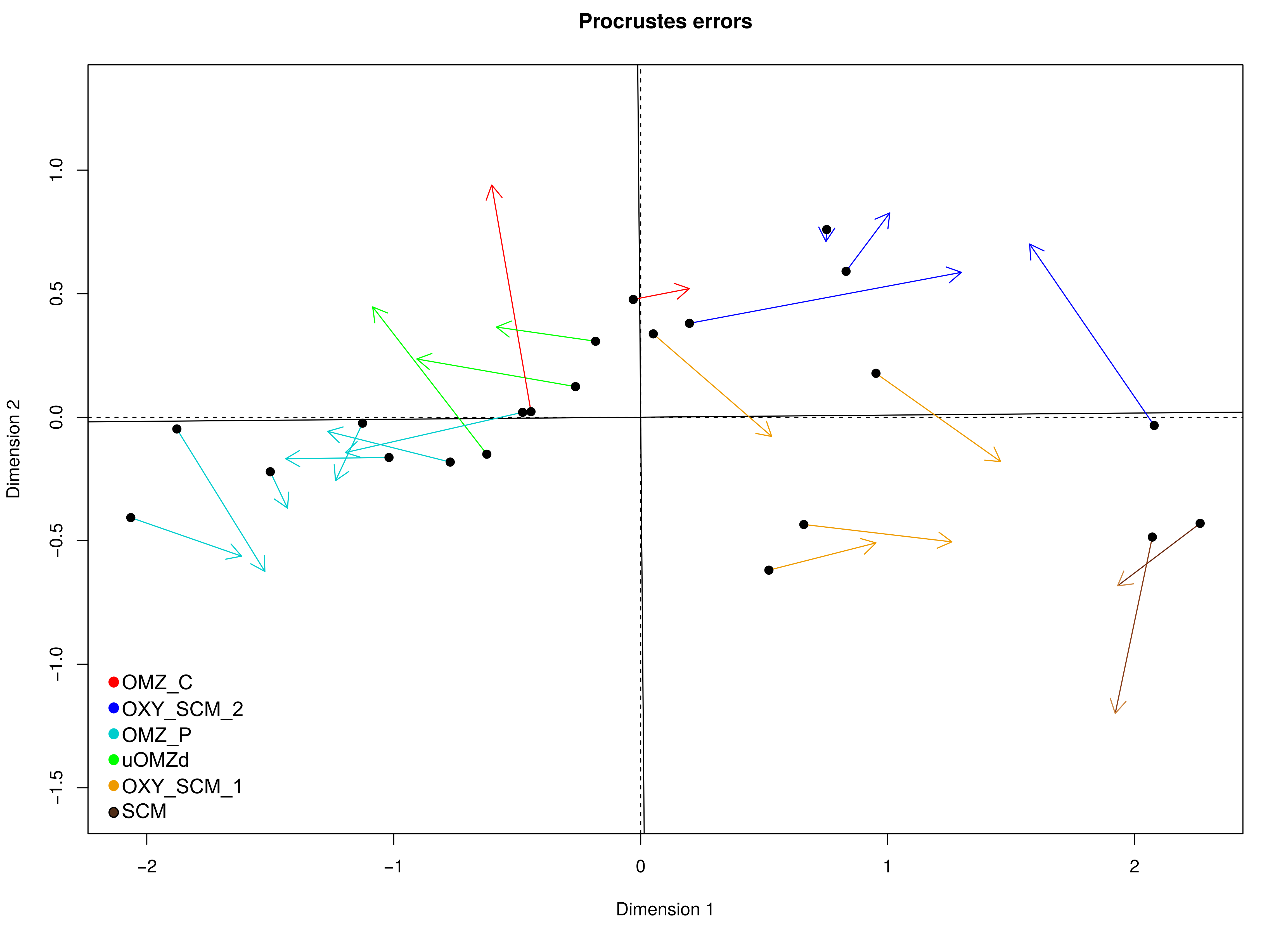

### Figures S7

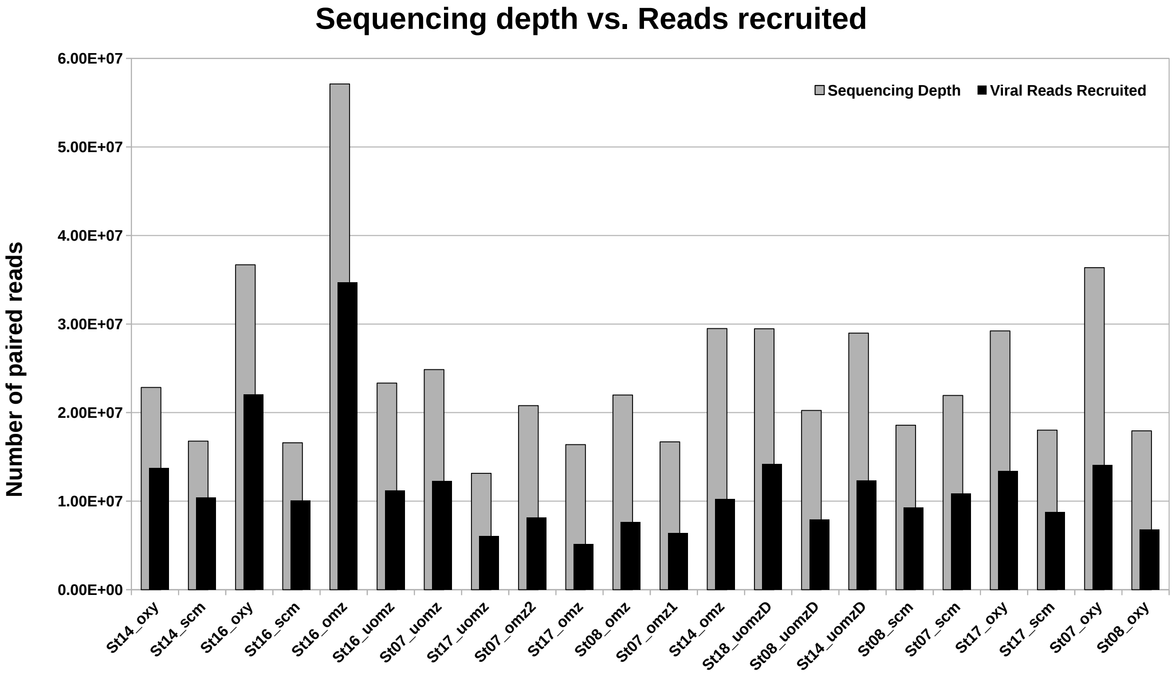
