## Supplementary material for "Genome-resolved viral ecology in a marine oxygen minimum zone (OMZ)": Figure S3

### APResult object

**Number of samples = 22**

**Number of iterations = 142**

**Input preference = -3303.617**

**Sum of similarities = -16882.67**

**Sum of preferences = -16518.08**

**Net similarity = -33400.76**

**Number of clusters = 5**

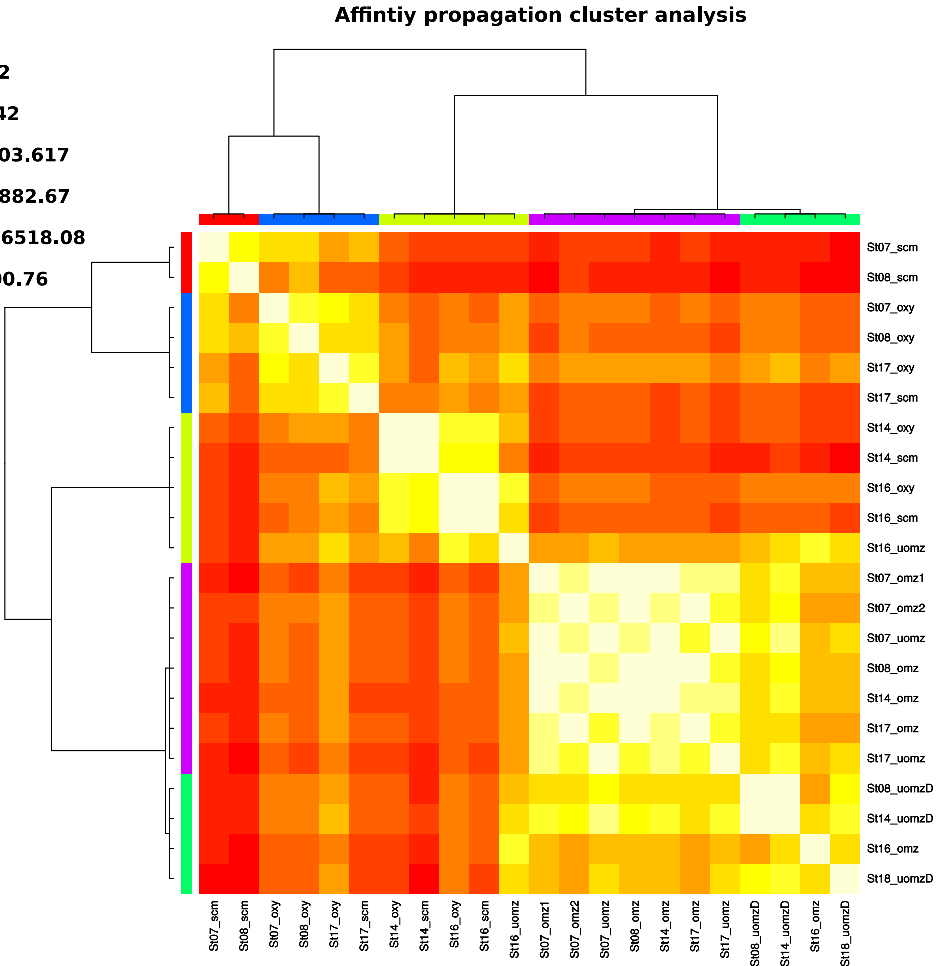
